# DETECTing Merkel cell Polyomavirus in Merkel Tumours

**DOI:** 10.1101/770537

**Authors:** Reety Arora, Komal Gupta, Anjali Vijaykumar, Sudhir Krishna

## Abstract

Merkel cell carcinoma (MCC) is a rare, aggressive skin cancer caused either by Merkel cell polyomavirus (MCV) T antigen expression, post integration (∼80% cases), or by UV mediated DNA damage. Interestingly, overall survival of patients suffering from MCV positive Merkel cell carcinoma is better, making this differential information of significant diagnostic and prognostic value. Also, MCV as a causative agent also provides a direct target for therapy in virus positive MCC patients. Currently, the methods used for diagnosis of MCV in tumours are often tedious, discordant and unreliable. In this study we used a guided molecular scissors based - DNA Endonuclease Targeted CRISPR Trans Reporter (DETECTR) technique to develop an *in vitro* molecular diagnostic tool for MCV positive MCC. DETECTR couples recombinase polymerase based amplification of target MCV DNA with Cas12a mediated detection. CRISPR diagnostics couple specific detection followed by cutting of the pathogenic DNA by the Cas enzyme – gRNA complex, with non-specific cutting of ssDNA that provides a measurable visual cue. To detect MCV DNA in MCC tumours, we designed Cas12a gRNAs targeting the MCV DNA and tested their targeting efficiency, and sensitivity using a fluorophore quencher labeled reporter assay. We show that this sophisticated MCV DETECTR system can detect MCV integrated in Merkel tumour rapidly, specifically and with femto-molar sensitivity. This new MCV DNA detecting system is promising and we hope it can be coupled with histopathological and immunohistochemical studies to diagnose the viral status of MCC in clinics in the near future.

## Introduction

Merkel Cell Carcinoma (MCC) is a rare and aggressive neuroendocrine skin cancer^1^. MCC is associated with old age, excessive UV exposure and weak immune system^2, 3^. It is either caused by Merkel cell polyomavirus (MCV)^4^ or long term Ultraviolet (UV) exposure^5-7^.

MCV is the latest addition to the group of human oncoviruses and the only known oncovirus in the polyomavirus family^4^. The virus has a double stranded DNA genome, encoding for early and late region genes. Early region genes encode for Small and Large Tumour antigens (sT and LT, respectively) and late region genes encode for the viral proteins (VP)^3, 4^.

In virus positive cancers MCV-dependent carcinogenesis requires two events. First, the viral genome gets integrated into the host cell genome, followed by LT getting truncation mutations in its DNA binding and helicase domains to make the virus replication incompetent^8^. The expressed T antigens then lead to oncogenesis by altering several pathways and are required for proliferation and survival of MCC cells^9-11^. A majority of MCC cases have been found to be MCV positive (∼80%)^1, 3, 4, 12^.

Both viral positive and viral negative Merkel tumours show identical phenotypes, immunohistochemical (IHC) staining patterns and occur at similar sun-exposed locations on the human body^3, 13-17^. A pathologist is unable to distinguish between the 2 types of tumours without the aid of Next-Gen-Sequencing^5, 18^. In a recent study, Moshiri S. et al. showed that out of 282 MCC tumours, 20% were MCV negative using multimodal qPCR and immunohistochemistry analysis^17^. Interestingly, the study indicated that as compared to MCV negative MCC patients, MCV positive MCC patients had significantly better progression-free survival, MCC-specific survival, and overall survival from MCC. Hence identifying the viral status is not only diagnostically (as the cancer cause) important, but important for prognostic predictability as well.

Besides diagnosis and prognosis, the knowledge of MCV integration and T antigen expression in MCC serves are a promising piece of information for MCC treatment as well. The viral DNA sequences and proteins serve as significant external targets that can be exploited for directed therapy for MCV positive cases.

Currently, most commonly used techniques for detecting MCV in MCC tumours include amplifying MCV DNA from DNA isolated from MCC tumours using PCR or immunohistochemistry (IHC) staining for MCV LT using monoclonal antibodies^1, 3, 15, 17^. However, these techniques are not the most concordant and reliable. For instance, the Moshiri study using 282 MCC samples used 3 techniques, IHC with CM2B4 and Ab3 and PCR, for identifying the viral status^17^. However, only 167 / 282 cases showed same viral status in all the three techniques whereas of 282 total cases, 199 cases were positive by qPCR, 205 by CM2B4-IHC and 254 by Ab3-IHC. Besides, these techniques are usually tedious and required trained personnel to perform them. Thus, there is a need for developing an accurate, sensitive and quick system for distinguishing viral negative vs viral positive MCC cases^3, 19-22^. In this study we developed a CRISPR/Cas12a based *in vitro* molecular diagnostic system for detecting MCV.

CRISPR (Clustered Regularly Interspaced Short Palindromic Repeats)/Cas system is a genome editing technology derived from the bacterial immune system^23^. CRISPR uses a guide RNA (gRNA) molecule that targets a Cas endonuclease to a specific genomic site using sequence homology and PAM (Protospacer adjacent motif) recognition. Upon target binding Cas protein induces a double strand break in the target^24-26^. Cas12a (Also known as Cpf1) is a type V CRISPR protein having various properties distinct from Cas9. Cas12a enzymes recognize a T nucleotide–rich protospacer-adjacent motif (PAM), catalyze their own guide CRISPR RNA (crRNA) maturation, and generate a PAM-distal dsDNA break with staggered 5′ and 3′ ends^23, 27, 28^. Interestingly, unlike Cas9, after dsDNA target binding the Cas12a enzyme leads to indiscriminate trans ssDNA cleavage activity^29^. Chen et. al. used this property to develop DNA endonuclease-targeted CRISPR trans reporter (DETECTR) system which can rapidly and specifically detect target HPV DNA^29^.

We adapted the DETECTR system to detect the presence of MCV DNA integration in the Merkel cell tumour genome. Briefly, we assemble a reaction mix containing AsCas12a, MCV specific gRNA, test DNA and fluorophore-quencher (FQ) ssDNA substrate in a tube. In the presence of MCV DNA, Cas12a binds and cleaves the cis target DNA followed by trans cleavage of fluorescently labelled ssDNA. This Cas12a mediated DNase activity leads to fluorescence based detection of MCV.

Here we show that when combined with Recombinase Polymerase based amplification of target DNA, DETECTR system can identify MCV with femtomolar sensitivity. DETECTR can detect MCV in MCV positive MCC cells efficiently and specifically. Thus, we hope that this MCV DNA detecting system can be coupled with histopathological and immunohistochemical studies to diagnose the viral status in MCC and help guide clinicians in the near future.

## Materials and Methods

### 1. sgRNA design

10 AsCpf1 gRNAs were designed to target the Non-coding Control Region (NCCR), small and large Tumour (sT and LT, respectively) antigen of MCV. Two CRISPR gRNA design tools: Benchling (https://benchling.com/pub/cpf1) and RGEN (www.rgenome.net/cas-designer) were used. MKL-1 (Genbank Accession #: FJ173815.1) and MS-1 (Genbank Accession #: JX045709.1) MCV sequences were used as target DNA sequences. gRNAs with highest specificity score and with lowest possible off targets were selected. MCV gRNA target region sequence conservation was analyzed using NCBI BLAST (https://blast.ncbi.nlm.nih.gov/Blast.cgi).

### 2. Synthesis of sgRNAs using IVT

For *in vitro* transcription (IVT), two oligonucleotides were designed as DNA templates, per gRNA. Forward oligonucleotide for AsCas12a gRNA synthesis consisted of the T7 promoter (TAATACGACTCACTATAGG) followed by AsCas12a sgRNA scaffold and 20-nucleotide long guide RNA. Three G’s were added after T7 promoter sequence for efficient T7 transcription. Target substrate for AsCas12a NCCR gRNA1 *in vitro* cleavage (pam/<u>TARGET</u>): 5’ GGCCTCTCTCTTTTtttc<u>CAGAGGCCTCGGAGGCTAGG</u>AGCCCCAAGCCTCTG 3’ crRNA oligo for *in vitro* transcription (**T7** / ADDED G / *SCAFFOLD* / <u>GUIDE</u>):

Forward oligo:

**TAATACGACTCACTATA**GGG*TAATTTCTACTCTTGTAGAT*<u>CAGAGGCCTCGGAGGCT</u> <u>AGG</u>

Reverse oligo:

<u>CCTAGCCTCCGAGGCCTCTG</u>*ATCTACAAGAGTAGAAATTA*CCC**TATAGTGAGTCGTA TTA**

Forward and reverse oligonucleotides were annealed by heating to 95°C for 5 minutes and slow cooling on bench top. sgRNAs were synthesized using the MEGAShortScript™ Kit (# AM1354, Invitrogen) following the manufacturer’s protocol with 2ug annealed oligonucleotides template, overnight at 37°C. The RNA was treated with 1ul of DNase TURBO for 15 minutes followed by purification using the MEGAclear Transcription Clean-Up Kit (# AM1908, Invitrogen).

### 3. *In vitro* cleavage assay

*In vitro* cleavage reaction was performed at 37°C in cleavage buffer consisting of 20 mM HEPES (pH 7.5), 150 mM KCl, 10 mM MgCl_2_, 1% glycerol and 0.5mM DTT. 30nM AsCas12a (#1081068, IDT) and 36nM gRNA were pre-assembled in the cleavage buffer at 37°C for 10 minutes. 18.5nM dsDNA target was added to the reaction (20ul). PCR amplicons of NCCR, sT and LT regions from plasmid RAZ2 (Addgene #114382) were used as dsDNA target templates. The PCR amplicon was purified using Ampure beads (#A63881, Beckman Coulter) as per manufacturer’s protocol. For M13 cleavage assays, 30nM AsCas12 and 36nM gRNA were preassembled at 37°C for 10 minutes in cleavage buffer. 40nM dsDNA activator and 10nM single stranded M13mp18 phage (#N4040S, NEB) were added to initiate the reaction (30ul). The reaction was incubated at 37°C for 60 minutes. The IVC reactions were stopped by treatment with 2ul of Proteinase K (10mg/ml) at 55°C for 10 minutes. 1x gel loading dye was added to the reactions and samples were run on 2% agarose gels (# RM273, HIMEDIA).

### 4. Fluorophore quencher (FQ)-labeled reporter assays

200nM AsCas12a, 250nM gRNA and 18.5nM dsDNA target template were pre-assembled in a 5ul reaction for 30 minutes at 37°C. Reaction was initiated by diluting these complexes to AsCas12a: gRNA: dsDNA template to 50nM: 62.5nM: 4.6nM in 1x Binding buffer (20 mM Tris-HCl, pH 7.5, 100 mM KCl, 5 mM MgCl2, 1 mM DTT, 5% glycerol, 50 µg ml^-1^ heparin) and 50nM custom ssDNA FQ reporter substrate (IDT). The custom ssDNA reporter substrate consisted of 5’ 6-FAM^™^, 5 nucleotides long oligo (TTATT) and 3’ Iowa Black® FQ. The 20ul reactions were incubated at 37°C for 1 hour in a 384 well microplate format. The fluorescence was measured using Tecan (Infinite M Plex) (Excitation: 485nm, Emission: 535nm). Experiments were repeated independently 3 times with 2 technical repeats each time. The graphs were drawn using GraphPad Prism and statistical significance was calculated using one way ANOVA.

### 5. Cell culture

HEK293T, U20S and BJhTERT cell lines were obtained from ATCC and grown in Dulbecco’s Modified Eagle Medium (DMEM) supplemented with 10% Fetal Bovine Serum (FBS), 1X Penicillin/Streptomycin (Pen Strep, catalog # 15140122). Merkel Cell Carcinoma cell line MKL-1 was obtained from ECACC (# 09111801) and MS-1, MCC26 and WaGa cells were kind gifts from Dr. James Decaprio’s laboratory at Dana Farber Cancer Institute, Boston. MCC cell lines were cultured in Roswell Park Memorial Institute (RPMI) Media with 10% FBS and 1X Penicillin/Streptomycin. The cells were grown at 37°C with 5% CO_2_.

### 6. Genomic DNA isolation

10^6^ cells were harvested for genomic DNA extraction. Cells were lysed in Tris EDTA buffer and 200ug/mL Proteinase K at 55°C for 1 hour. After Proteinase K was heat inactivated, 1ul of this lysed sample was used for DETECTR experiments.

### 7. Recombinase Polymerase Reaction (RPA)

For DETECTR experiments, genomic DNA was first amplified using recombinase polymerase reaction (RPA). The RPA reaction mix was prepared by adding 0.48uM forward primer and reverse primers, 29.5 µl primer Free Rehydration buffer, 1µL of extracted genomic DNA and water to 13.2 µl (Total volume 47.5 µl). The reaction mix was added to one TwistAmp® Basic (#TABAS03KIT, TwistDx) reaction tube. 2.5µl of 280mM Magnesium Acetate (MgOAc) was added and mixed to start the reaction. The reaction was incubated at 37°C for 20 minutes.

### 8. DETECTR assay for genomic DNA

DETECTR uses RPA reaction to amplify target DNA followed by Cas12a based detection. 18µL of the RPA reaction mix was assembled with 2µL of 50nM AsCas12a, 62.5nM gRNA and 50nM custom ssDNA FQ substrate. Reaction was incubated at 37°C for 1 hour and fluorescence was measured using Tecan Infinite Pro 200.

For investigating DETECTR’s sensitivity, a titration experiment was performed. NCCR, sT and LT target region was amplified from plasmid RAZ2 (Addgene #114382) and used as dsDNA target. This dsDNA was diluted to varying concentrations ranging from 10^−8^ to 10^−15^M. Fluorophore quencher (FQ)-labeled reporter assay was performed as described previously using these targets with or without RPA.

## Results

### gRNAs were designed to target all MCV variants

To detect the presence of integrated MCV DNA in MCC cells, we designed 10 guide RNAs compatible with AsCas12a. Three gRNAs are complementary to the Non-Coding Control Region (NCCR), 5 gRNAs target Exon 1 of T antigen and the remaining 2 gRNAs target Exon 2. We used MKL-1 MCV sequence (Genbank Accession #: FJ173815.1), till the conserved Retinoblastoma (Rb) binding domain region to design these gRNAs (Figure 1 and Table 1). All gRNAs designed targeted conserved regions across MCV subtypes known so far ^30-32^. Using Clustal Omega Multiple Sequence Alignment we checked 52 MCV sequences (from NCBI) across all gRNA-targeting regions.

**Table 1.**
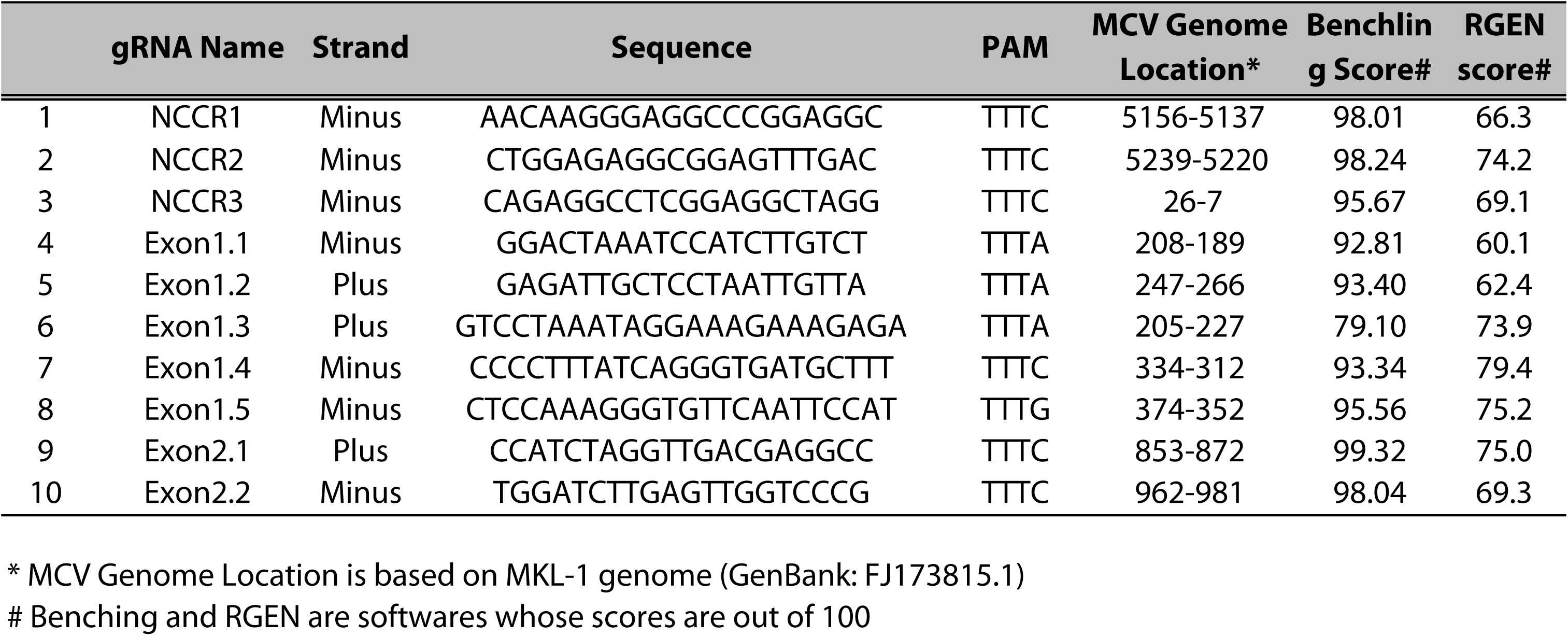
AsCas12a_gRNAs against Merkel Cell Polyomavirus.

**Figure 1:**
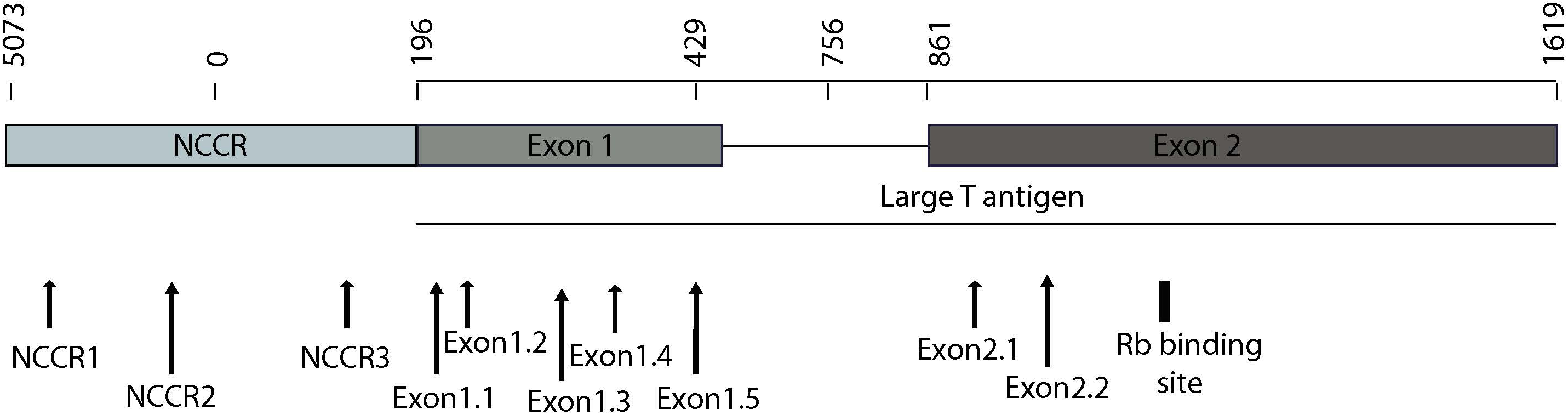
MCV NCCR And Early Region Map Showing gRNA Locations. The map above shows the location of 10 gRNAs (AsCas12a CRISPR based) that we designed and the specific sites on the MCV genome that they target.

### Screening and validation of gRNAs targeting MCV genome

To access the cleavage efficiency of these gRNAs, we performed *in vitro* cleavage assays. We used the 2193 bp long NCCR and T antigen region (derived from MS-1) as template/ DNA substrate for our assays. Six out of 10 gRNAs efficiently cleaved the target substrate as shown by the cleaved bands (Figure 2A).

**Figure 2:**
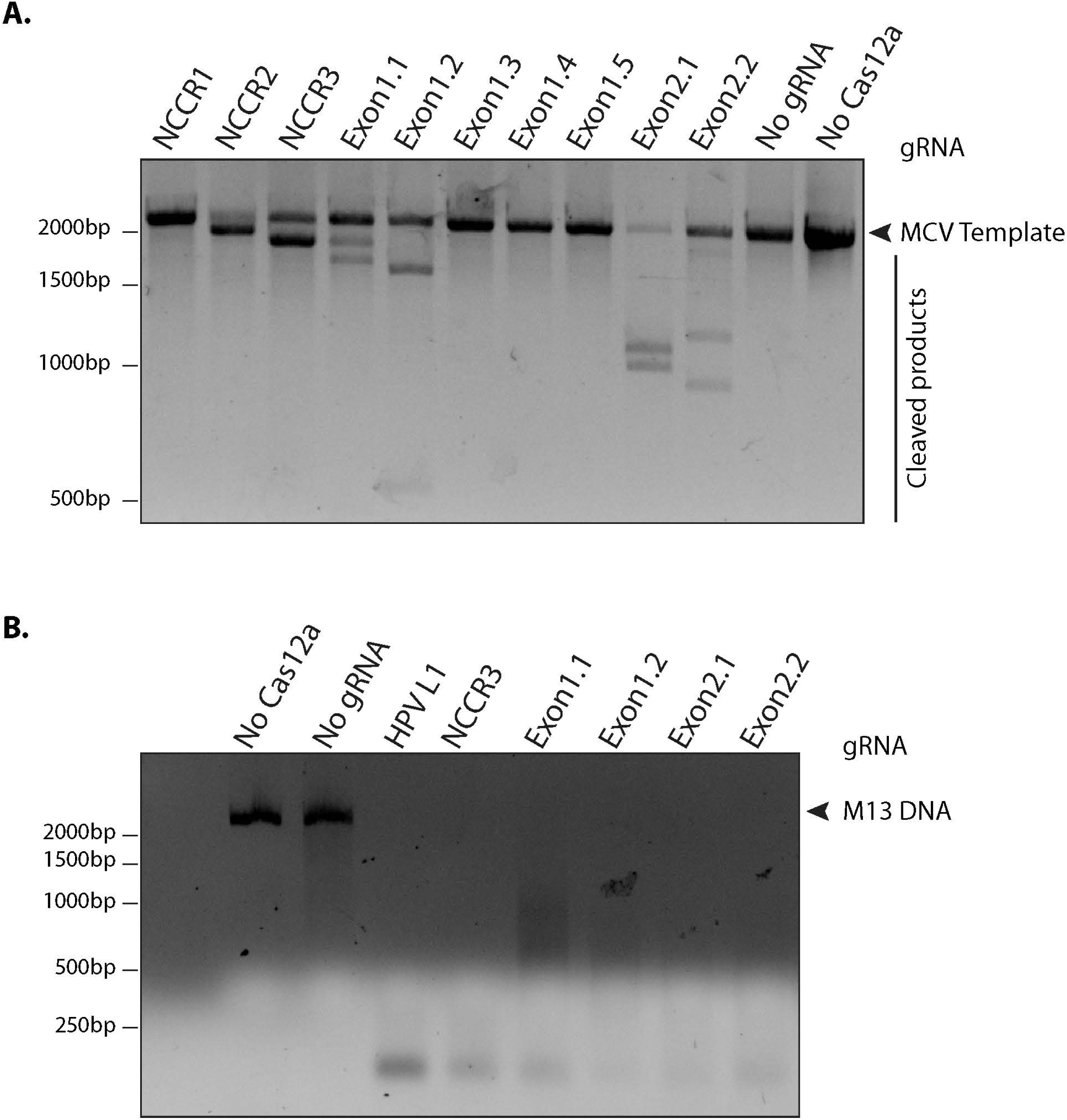
Screening gRNAs For Efficient Cleavage. **(A). *In vitro* Cleavage (IVC) Assay For All gRNAs.** An MCV Template of 2193 bp spanning the NCCR region to the RB-binding site of T antigen was used for the assay (shown by the arrow). Of the 10 gRNAs tested, six showed efficient cleavage that resulted in visible chopped bands on a 2% agarose gel. **(B.) Non-specific Trans ss DNA Shredding Activity Of AsCas12a-gRNA.** M13 DNA phage reporter was subjected to MCV gRNA, complementary ssDNA target (cis-activator) and AsCas12a. All 5 gRNAs, namely NCCR3, Exon1.1, Exon1.2, Exon2.1 and Exon2.2 resulted in indiscriminate shredding of single-stranded M13 DNA.

Next, to validate the non-specific trans ssDNA shredding activity of AsCas12a, we used a non-complementary, circular, single stranded M13 DNA phage reporter. In the presence of MCV gRNA and complementary ssDNA target (cis - activator), AsCas12a led to indiscriminate shredding of single stranded M13 phage (Figure 2B). Thus, 5 MCV gRNAs: NCCR3, Exon 1.1, Exon 1.2, Exon 2.1 and Exon 2.2, were selected for further studies for detecting MCV.

### Detection of MCV via fluorescence measurement

To develop a MCV DNA detection system, we used a fluorophore quencher (FQ)–labeled reporter assay. We assembled AsCas12a with its MCV gRNA and a complementary dsDNA cis target. We introduced a non-specific ssDNA FQ reporter to this reaction. In the presence of gRNA targeting MCV DNA and complementary target MCV DNA, AsCas12a cleaved the ssDNA FQ reporter as observed through the emitted fluorescence. We found that all the 5 selected gRNAs showed significant fold change in emitted fluorescence as compared to no gRNA reaction (Figure 3 and Supplementary Figure 1). HPV L1 targeting gRNA and matching template were used as a positive control for the assays (Supplementary Figure 2)^29^.

**Figure 3:**
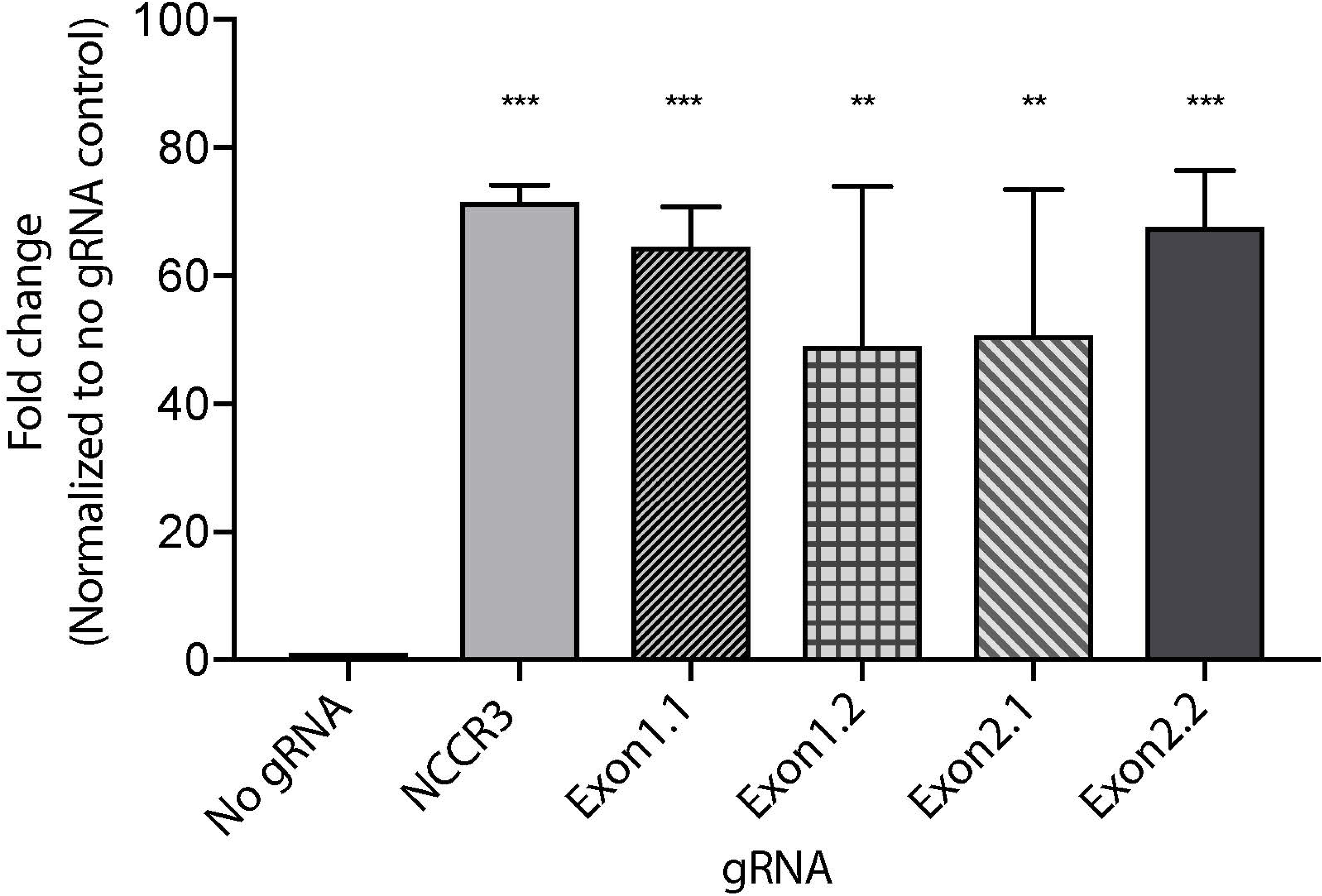
MCV Detection Via Fluorescence Measurement. A Fluorophore-Quencher (FQ) labeled reporter assay was used to test the gRNA-AsCas12a combinations. MCV gRNA, AsCas12a and complementary dsDNA cis target were assembled and subjected to a custom ssDNA FQ reporter (excitation 485nm, emission 535 nm). All 5 gRNAs showed significant fold change in emitted fluorescence as compared to No gRNA control (∼48 fold and above). Error bars represent SD for three independent experiments. One-way ANOVA with Dunnett test was performed for statistical analysis. (Adjusted p values: p_NCCR_=0.0003, p_Exon1.1_=0.0007, p_Exon1.2_=0.0065, p_Exon2.1_=0.005 and p_Exon2.2_=0.0005)

We proceeded further with 2 gRNAs: NCCR 3 and Exon 1.1. The DETECTR from Prof. Doudna’s group, couples Recombinase Polymerase with Cas12a based DNA detection ^29^. To investigate the sensitivity of our MCV DETECTR system, we used a plasmid containing the MS-1 NCCR and T antigen region with varying concentrations (range-10^−15^ to 10^−8^ Molar). We performed Cas12a based detection using these targets with and without Recombinase Polymerase Amplification. Our results show that the MCV DETECTR (RPA + Cas12a) system we created can efficiently detect MCV DNA up to femtomolar concentration (Figure 4 and Supplementary Fig 3).

**Figure 4:**
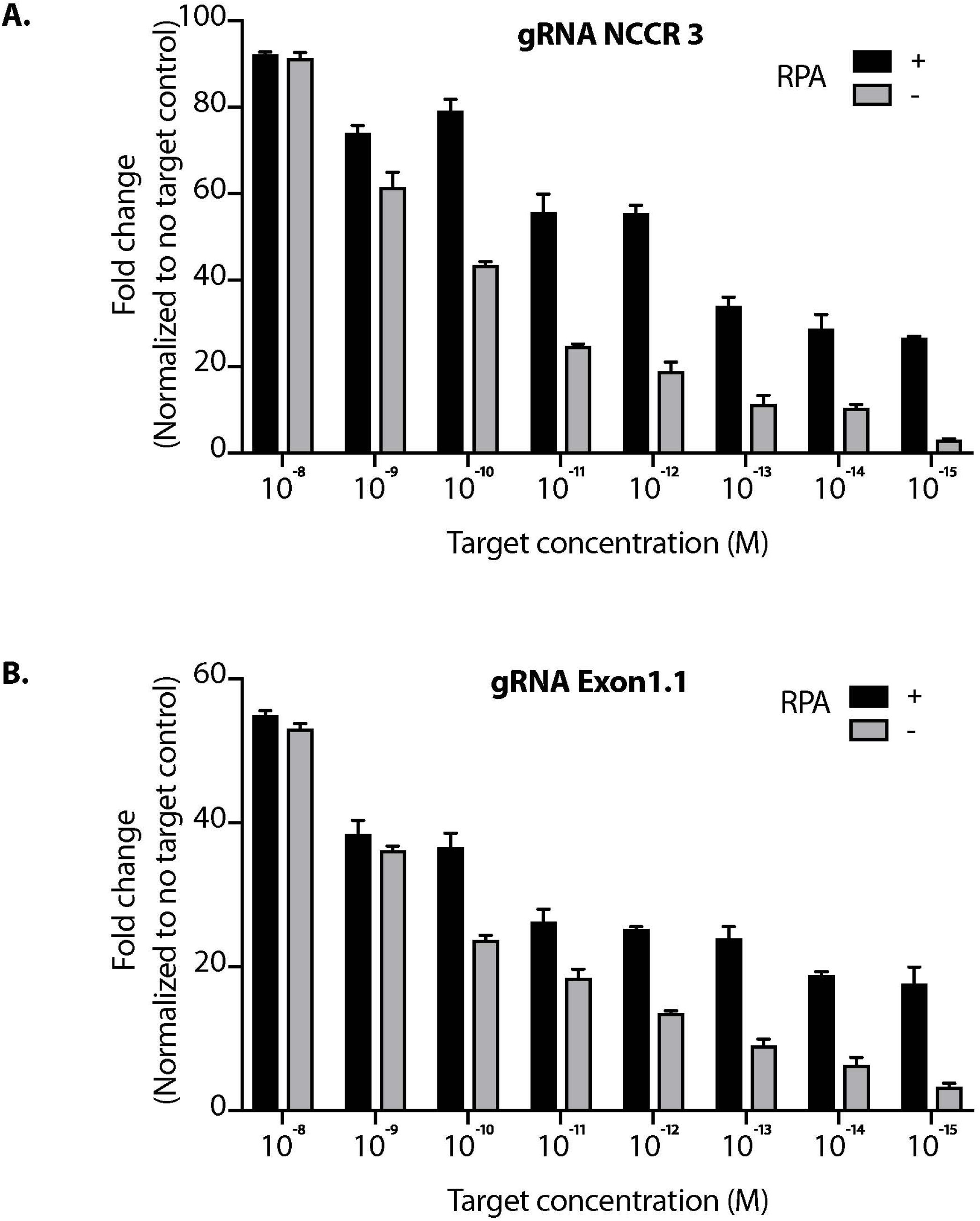
Sensitivity Of MCV gRNA To Detect Low Copies of MCV. The Cas12a based detection of MCV was performed in the presence and absence of RPA (Recombinant Polymerase Amplification) with varying concentrations (range-10^−15^ to 10^−8^ M target concentration) for both (A.) gRNA NCCR3 and (B.) gRNA Exon1.1. Error bars represent SD for three independent experiments.

### DETECTR can diagnose MCV in MCC genomic DNA

Next, we investigated whether DETECTR can specifically and efficiently detect MCV in MCV positive MCC cell’s genomic DNA. We extracted genomic DNA from various cultured human cells infected with MCV (MKL-1, MS-1, WaGa) or without MCV (MCC26, BJhTERT, U2OS). Whereas MKL-1, MS-1, WaGa and MCC26 are all Merkel cell carcinoma cell lines, BJhTERT is an immortalized fibroblast cell line and U2OS is an osteosarcoma cell line.

We performed MCV detection in these samples using AsCas12a DETECTR (RPA + Cas12a). We found that although only Cas12a-gRNA complex was not sensitive enough to identify MCV, DETECTR unambiguously detected MCV in MKL-1, MS-1 and WaGa cells (Figure 5). Also, there was no significant signal in the presence of non-complimentary DNA from MCV negative cells. Thus, we show that the MCV DETECTR can be potentially used as a specific and efficient platform for diagnosing MCV in MCC tumours (Figure 6).

**Figure 5:**
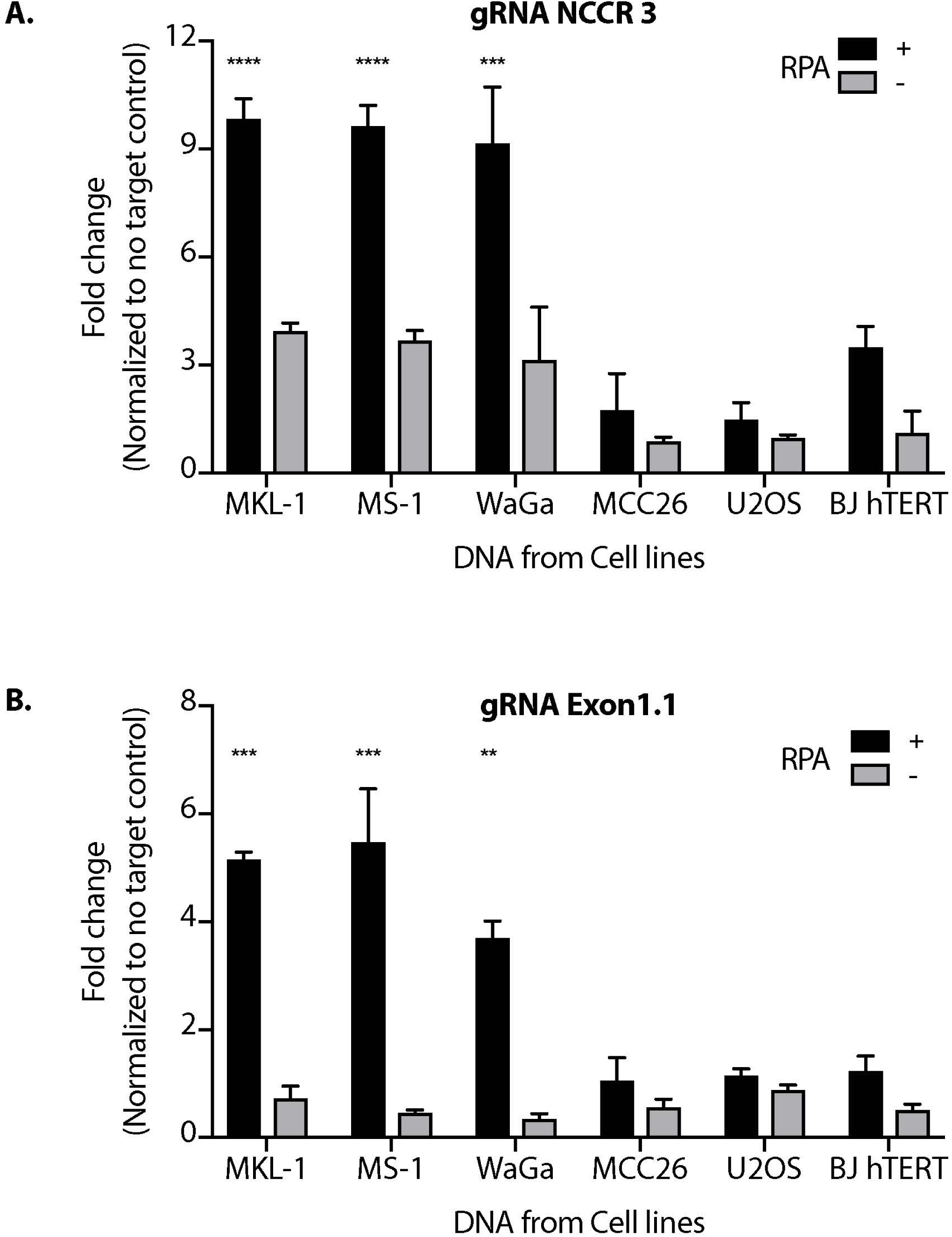
MCV DETECTR In Action. Genomic DNA from MCV positive MCC cell lines MKL-1, MS-1, WaGa; MCV negative MCC cell lines MCC26; Osteosarcoma cell lines U2OS and immortalized Fibroblasts BJhTERT were extracted and subjected to the MCV DETECTR assay. With the use of RPA, MCV was detected significantly in all MCV MCC positive cell lines for both (A.) gRNA NCCR and (B.) gRNA Exon 1.1. Fluorescence was normalized to no-target control and fold change plotted. Error bars represent SD for two independent experiments Dunnett test was performed for statistical analysis. (***= pvalue=0.001, **** = pvalue <0.0001).

**Figure 6:**
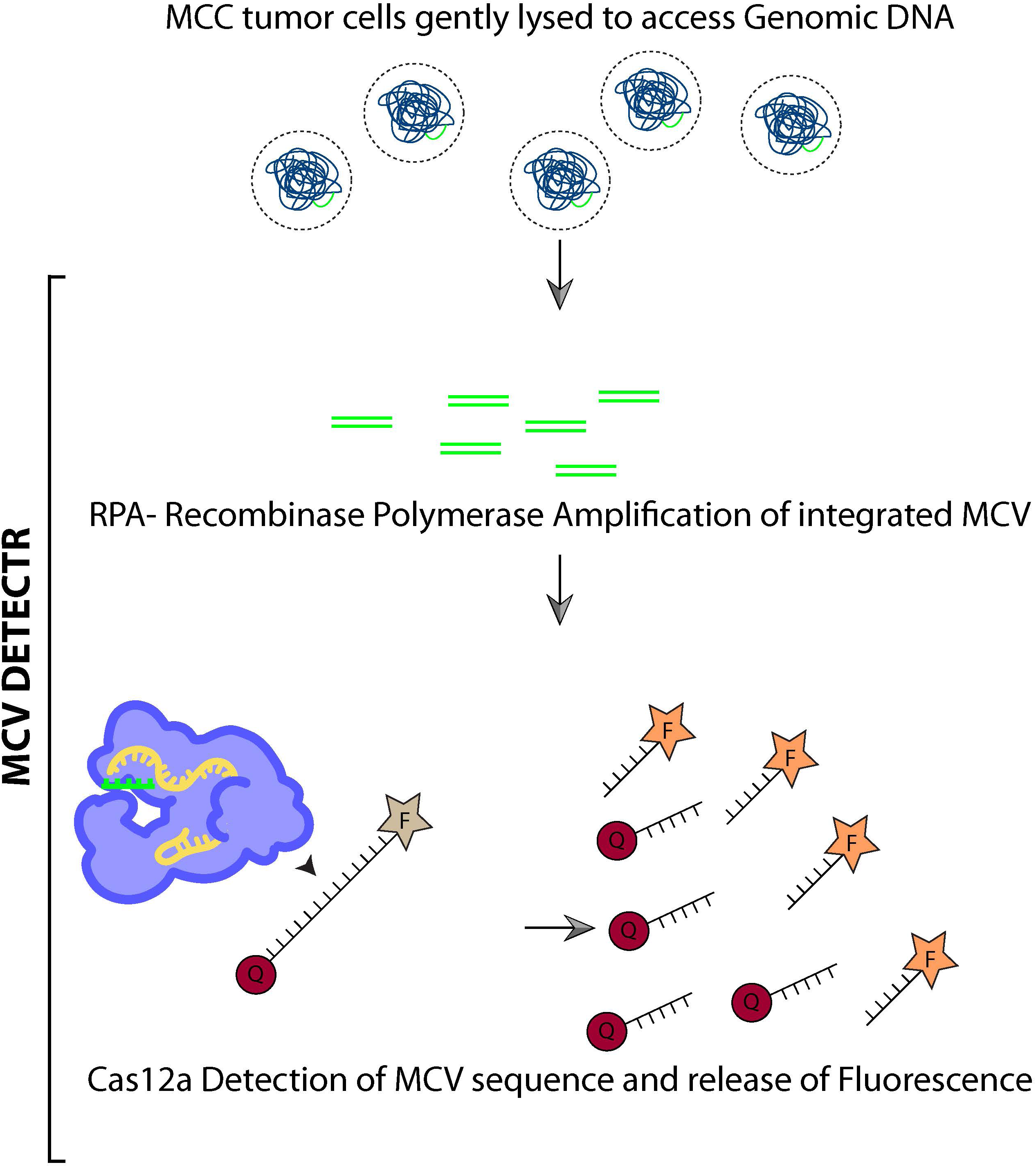
Schematic summarizing MCV DETECTR.

## Discussion

Currently, the methods used for the experimental detection or diagnosis of MCV in tumours include immunohistochemistry, PCR, RNA or DNA in situ hybridization and next-generation sequencing (NGS)^3, 12, 33-37^. All of these assays vary considerably in terms of sensitivity and specificity for MCV detection in tumours. Immunohistochemical measurement of T antigen protein expression is the commonest approach for MCV detection and the antibody used for the same is usually CM2B4 (88% sensitivity, 94% specificity)^17^. Recently Moshiri et al. introduced a more efficient multimodal approach combining PCR and immunohistochemistry for detection of viral T antigens^17^.

Although sensitive, these methods do have their drawbacks. For examples, CM2B4 often gives non-specific staining in both tonsillar and lymphoid tissues^17, 22^. Other antibodies that detect MCV T antigens, such as Ab3 and Ab5 have been reported, however none have been used as frequently as CM2B4 for detecting viral proteins in the clinic^3, 38^.

PCR is another easy and common method used for MCV detection by many laboratories. In comparison with the multimodal approach, PCR-based amplification within the second exon of LT had 83% sensitivity and 81% specificity^17^. Quantitative PCR (qPCR) extends this analysis and allows for estimation of number of integrated MCPyV copies per host cell genome^39, 40^. MCPyV copy number estimates have been reported to range from <1 copy per 100 cells to thousands of copies per cell^12, 36^.

Technical factors, including inefficient PCR amplification owing to mutations in the integrated MCPyV genome, primer incompatibilities, low purity of tumour samples or the detection of infectious wild-type MCPyV in the adjacent nonmalignant skin often confound the results for such tests ^19, 35^. Besides, qPCR does not allow for visual confirmation that positive results are associated with tumour cells; hence, background MCV presence cannot be excluded in MCC tumours with low signal^35^.

RNA in situ hybridization and NGS are other newer approaches being investigated for MCPyV detection. RNA in situ hybridization might provide PCR level sensitivity in conjunction with visual correlation with tissue morphology and the exclusion of background infection^35^. NGS, on the other hand is effective in detecting MCPyV sequences, including tumour-specific truncating mutations and viral integration sites^7, 33^. However, the time, expense and expertise required for these methods currently make these approaches impractical for many diagnostic and research laboratories.

The “cut and paste” function of CRISPR is popular and what the system is often associated with, after a guide RNA finds a target DNA sequence, the Cas nuclease makes a double-stranded cut and one can paste a sequence of choice at that site. The newer Cas12 and Cas13 have enabled researchers to use the “search” function of CRISPR in diagnostics^41^. Platforms such as DETECTR by Jennifer Doudna’s group^29^ and SHERLOCK^42, 43^ by Feng Zhang’s group have leveraged this search functionality to get sensitive and specific diagnostic results for HPV and Dengue virus respectively. Further studies to use CRISPR for Zika virus, Lassa fever and even Ebola are underway^44-47^. The startups called Mammoth Biosciences and SHERLOCK Biosciences are such endeavors for providing better diagnostics for disease.

These non-editing applications of CRISPR have brought to light a precise, easy and efficient detection system for disease and here we have extended Dr. Doudna’s group’s studies to the tumour virus Merkel cell polyomavirus.

Our work offers a novel diagnostic method for MCV detection (Figure 6). Our MCV DETECTR approach is accurate, efficient, easy to use and rapid (30 mins). It has a clear readout (fluorescence), is non-invasive, specific and sensitive to femto-molar levels. The simplicity of this method, lack of complex sample preparation and no requirement for expensive equipment makes it superior to others currently being used. This CRISPR trans reporter system for DNA detection has opened a new applications door for CRISPR based technologies with rapid and specific detection of Merkel cell polyomavirus in patient samples. Additionally, combining this approach with immunohistochemistry and/or PCR will substantially improve the sensitivities of detection of MCV.

Thereby, our system is a simple platform for molecularly diagnosing whether a tumour is virus positive or negative. This information is not only an important causal indicator for Merkel tumours but also a powerful prognosis predictor. We hope that it will soon be incorporated into clinical practice and positively affect MCC studies.

## Supporting information

All Supplementary Figures

## Acknowledgements

We are grateful to Dr. Debojyoti Chakraborty and Mohammed Azhar from IGIB, Delhi for guidance and advice on CRISPR related experiments. We thank Aurelie Jory for providing Cas12a protein for pilot experiments. Special thank you to Lamiya Dohadwala and Bhavana Nayer for technical assistance and to Prof Sanjeev Galande for scientific comments. We also thank Dr. Anirudha Lakshminarasimhan, Divij Kinger, Dr. Alpana De and Gayathri Rathinam from Tata Institute for Genetics and Society (TIGS), Bangalore for providing access and assistance in fluorescence measurements using Tecan machine. RA would also like to thank Zaina, Ziyah and Karan for their love, strength and support.

For Figure 6 the Innovative Genomics Institute (IGI) Glossary Icon Collection image for Cas12a was used. This is licenced under a Creative Commons Attribution-Non Commercial-ShareAlike 4.0 International Licence.

