## Supplementary material for "DETECTing Merkel cell Polyomavirus in Merkel Tumours": All Supplementary Figures

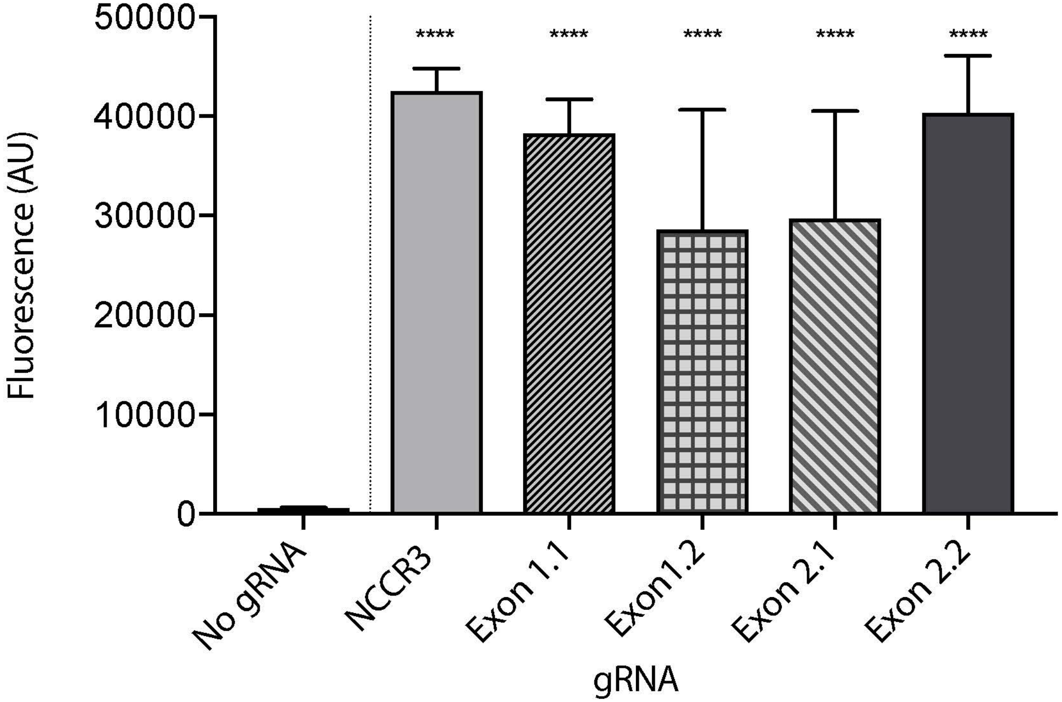


**Supplementary Figure 1. MCV Detection via Fluorescence Measurement**.
A Fluorophore-Quencher (FQ) labeled reporter assay was used to test the gRNA-AsCas12a combinations. MCV gRNA, AsCas12a and complementary dsDNA cis target were assembled and subjected to a custom ssDNA FQ reporter (excitation 485nm, emission 535 nm). All 5 gRNAs showed significant emitted fluorescence as compared to No gRNA control. Error bars represent SD for three independent experiments. One-way ANOVA with Dunnett test was performed for statistical analysis. ($p$ value $< 0.0001$)

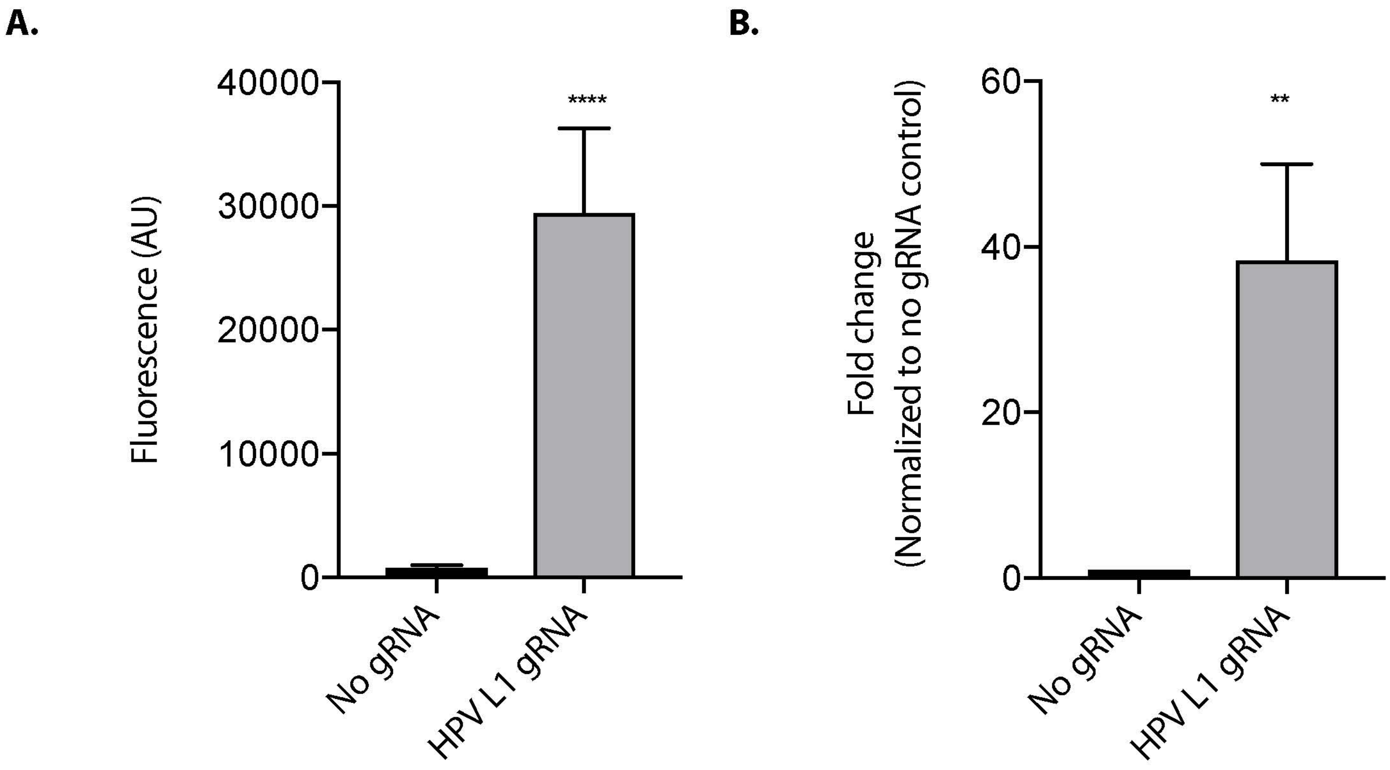


**Supplementary Figure 2. HPV L1 gRNA Fluorescence-Quencher (FQ)labeled Reporter Assay**

The Fluorophore-Quencher (FQ) labeled reporter assay that we designed was also used to test the HPVL1 gRNA previously reported by Chen. et al, 2018. HPV L1 gRNA, AsCas12a and complementary dsDNA cis target were assembled and subjected to a custom ssDNA FQ reporter (excitation 485nm, emission 535 nm). The gRNAs showed significant emitted fluorescence as compared to No gRNA control for both (A). FLuorescence and (B). Fold change plots. Error bars represent SD for three independent experiments. Two-tailed t test showed p value < 0.0001 (****) for fluorescence measuremnet and p value= 0.0051(**) when the measurement was normalized to No gRNA and plotted as fold change.

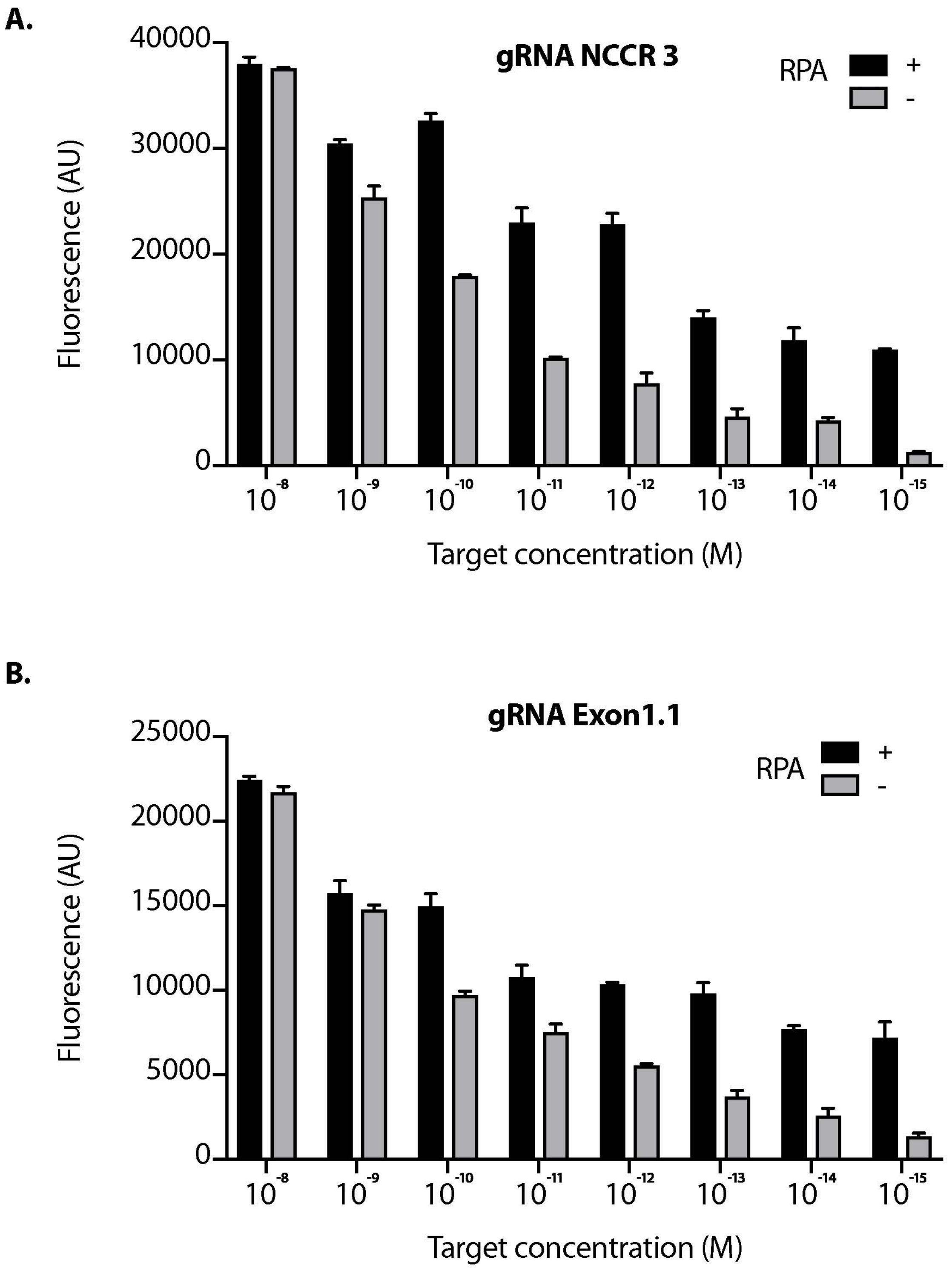


**Supplementary Figure 3. Sensitivity of MCV gRNA**
The Cas12a based detection of MCV was performed in the presence and absence of RPA (Recombinant Polymerase Amplification) with varying concentrations (range- femto M to 10 nM target) for both (A.) gRNA NCCR3 and (B.) gRNA Exon1.1.

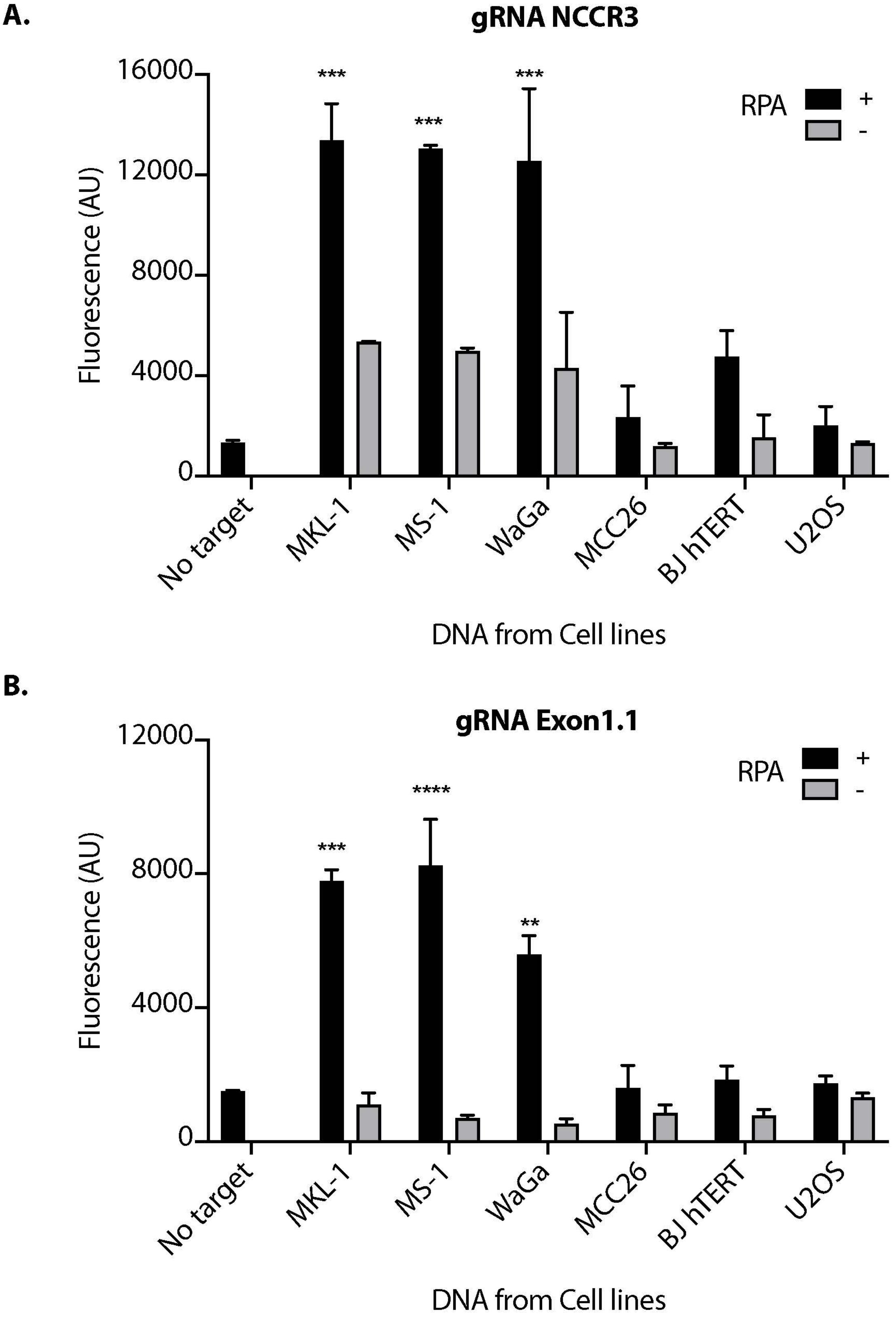


**Supplementary Figure 4. MCV DETECTR In Action (Fluorescence values)**
Genomic DNA from MCV positive MCC cell lines MKL-1, MS-1, WaGa; MCV negative MCC cell lines MCC26; Osteosarcoma cell lines U2OS and immortalized Fibroblasts BJhTERT were extracted and subjected to the MCV DETECTR assay. With the use of RPA, MCV was detected significantly in all MCV MCC positive cell lines for both (A.) gRNA NCCR ($p_{(MKL\text{-}1)} = 0.0003$, $p_{(MS\text{-}1)} = 0.0003$, $p_{(WaGa)} = 0.0004$) and (B.) gRNA Exon 1.1 ($p_{(MKL\text{-}1)} = 0.0001$, $p_{(MS\text{-}1)} < 0.0001$, $p_{(WaGa)} = 0.0018$).Error bars represent SD for two independent experiments Dunnett test was performed for statistical analysis.

**Supplementary Table 1: Primers**

| | | |
|---|---|---|
| **Oligonucleotides for AsCas12a gRNA in vitro synthesis** | | |
| 1 | HPV L1 gRNA F | TAATACGACTCACTATAGGGTAATTTCTACTCTTGTAGATCTACATTACAGGCTAACAAA |
| 2 | HPV L1 gRNA R | TTTGTTAGCCTGTAATGTAGATCTACAAGAGTAGAAATTACCCTATAGTGAGTCGTATTA |
| 3 | MCV.NCCR1.F | TAATACGACTCACTATAGGGTAATTTCTACTCTTGTAGATAACAAGGGAGGCCCGGAGGC |
| 4 | MCV.NCCR1.R | GCCTCCGGGCCTCCCTTGTTATCTACAAGAGTAGAAATTACCCTATAGTGAGTCGTATTA |
| 5 | MCV.NCCR2.F | TAATACGACTCACTATAGGGTAATTTCTACTCTTGTAGATCTGGAGAGGCGGAGTTTGAC |
| 6 | MCV.NCCR2.R | GTCAAACTCCGCCTCTCCAGATCTACAAGAGTAGAAATTACCCTATAGTGAGTCGTATTA |
| 7 | MCV.NCCR3.F | TAATACGACTCACTATAGGGTAATTTCTACTCTTGTAGATCAGAGGCCTCGGAGGCTAGG |
| 8 | MCV.NCCR.3.R | CCTAGCCTCCGAGGCCTCTGATCTACAAGAGTAGAAATTACCCTATAGTGAGTCGTATTA |
| 9 | MCV.Exon 1.1 F | TAATACGACTCACTATAGGGTAATTTCTACTCTTGTAGATGGACTAAATCCATCTTGTCT |
| 10 | Exon 1.1 R | AGACAAGATGGATTTAGTCCATCTACAAGAGTAGAAATTACCCTATAGTGAGTCGTATTA |
| 11 | MCV.Exon 1.2 F | TAATACGACTCACTATAGGGTAATTTCTACTCTTGTAGATGAGATTGCTCCTAATTGTTA |
| 12 | MCV.Exon 1.2 R | TAACAATTAGGAGCAATCTCATCTACAAGAGTAGAAATTACCCTATAGTGAGTCGTATTA |
| 13 | MCV.Exon 2.1 F | TAATACGACTCACTATAGGGTAATTTCTACTCTTGTAGATCCATCTAGGTTGACGAGGCC |
| 14 | MCV.Exon 2.1 R | GGCCTCGTCAACCTAGATGGATCTACAAGAGTAGAAATTACCCTATAGTGAGTCGTATTA |
| 15 | MCV.Exon 2.2 F | TAATACGACTCACTATAGGGTAATTTCTACTCTTGTAGATTGGATCTTGAGTTGGTCCCG |
| 16 | MCV.Exon 2.2 R | CGGGACCAACTCAAGATCCAATCTACAAGAGTAGAAATTACCCTATAGTGAGTCGTATTA |
| **Primers for RPA of MCV target** | | |
| 17 | NCCR3_RPA_FP | GCAGCAATAAAAGTTCAATCATGTAACCACAA |
| 18 | NCCR3_RPA_RP | CTTGGGATCTGCCCTTAGATACTGCCTTTTT |
| 19 | Exon1.1_RPA_FP | TAGTGAGGTAGCTCATTTGCTCCTCTGCTCTT |
| 20 | Exon1.1_RPA_RP | TTGCCATAACAATTAGGAGCAATCTCTAAAAG |
| 21 | Exon1.2_RPA_FP | CTCCTTCTGCATATAGACAAGATGGATTTAGTC |
| 22 | Exon1.2_RPA_RP | ATTCCATCATTATAACAGGATTTCCCCCTTTAT |
| 23 | Exon 2.1_RPA_FP | GTAAGTATTAGATATGGAAAAGTCTATAAGGCAA |
| 24 | Exon 2.1_RPA_RP | GATTCAGCTTCGGGAAGGCATACGAATATGG |
| 25 | Exon 2.2_RPA_FP | GGGACCACTAAATTCAAAGAATGGTGGAGATC |
| 26 | Exon 2.2_RPA_RP | ACGCTGAGAAGGACCCATACCCAGAGGAAGAG |
| 27 | HPV16_RPA_FP | TTGTTGGGGTAACCAACTATTTGTTACTGTT |
| 28 | HPV16_RPA_RP | CCTCCCCATGTCGTAGGTACTCTTTAAAG |
| **Primers for PCR amplifying Cas12a MCV IVC template** | | |
| 29 | HPV L1_IVC target_FP | GGTCGTGGTCAGCCATTAGG |
| 30 | HPV L1_IVC target_RP | GTGGTGGGTGTAGCTTTTCG |
| 31 | RAZ2_IVC target_FP | GACTACATCGATCCTGAAAAATAAATAAG |
| 32 | RAZ2_IVC target_RP | AGCACGATATCCTAATTGAGCAGTGGCTTATTCTCTTG |
